# Benchmarking biochemical networks generated by large language models

**DOI:** 10.1101/2025.07.28.667217

**Authors:** Jeevan Tewari, Benjamin W. Dahl, B. Adam Bates, Jason A. Papin, Jeffrey J. Saucerman

## Abstract

Computational models of biochemical networks provide frameworks for predicting how molecular cues guide cell decisions. These models are typically limited by the time-intensive manual curation required to extract network mechanisms from incomplete literature. Here, we test whether general-purpose large language models (LLMs) can generate accurate models of signaling and metabolic networks. We find that general-purpose LLMs generate 24-65% of the reactions of literature-curated signaling networks for cardiomyocyte hypertrophy, myofibroblast activation, and mechanosignaling. Further, logic-based models based on these networks predict responses to perturbations with accuracies of 6-33%. In the context of metabolic modeling, LLMs are able to generate 64-91% of the reactions within the core *Escherichia coli* metabolic network and demonstrate highly variable accuracies in predicting substrate utilization. Current general-purpose LLMs generate biochemical networks with moderate accuracy, and this study provides a pipeline and benchmarks to guide future improvements.

---

The ability of cells to grow or respond to external stimuli is fundamental to life, mediated by complex metabolic and signaling networks. Computational models of signaling networks have been widely used to understand their systems properties and guide new experiments^1^, but they typically require substantial manual literature curation of network reactions and rate constants^2^. Similarly, metabolic network models provide a powerful framework for predicting cellular metabolism and substrate utilization, but their construction has historically required manual curation of stoichiometric coefficients, reaction reversibility, and metabolite–reaction connectivity.

Artificial intelligence (AI), and large language models (LLMs) in particular, have demonstrated strong performance in tasks that traditionally required labor-intensive knowledge curation. For example, general-purpose LLMs have scored highly on the U.S. Medical Licensing Examination^3^, law exams^4^, and Advanced Placement exams^4^. AI has also made substantial contributions in biology, including predicting protein structure^5^ and cell types^6^, and is expected to play a central role in the development of comprehensive virtual cells^7,8^. However, to our knowledge, LLM-generated predictive models of biochemical networks have not been reported or benchmarked. Here, we develop pipelines for LLM-generated models of signaling and metabolic networks, and we test whether general-purpose LLMs accurately predict network structure. Additionally, we assess the LLM-generated networks’ accuracy in predicting perturbation responses, as benchmarked against manually curated models and experimental evidence^9–11^.

We first established a computational pipeline to generate models of signaling networks from LLMs using a specified gene set (**Figure 1A**). For benchmarking, each gene set was derived from annotations of three manually curated, logic-based network models that have been extensively validated using experimental data^9–11^. We optimized a programmatic prompt that iteratively queries an LLM for directed interactions (activation or inhibition) among the gene products in the context of a particular phenotype (here “hypertrophy”, “fibroblast”, or “mechanosignaling”). We then extracted the returned directed interactions, examined network structure, and converted the network into a logic-based formalism to simulate network responses to perturbations^12^. Each LLM was prompted 10 times per network to examine reproducibility (see methods for prompting parameters).

**Figure 1:**
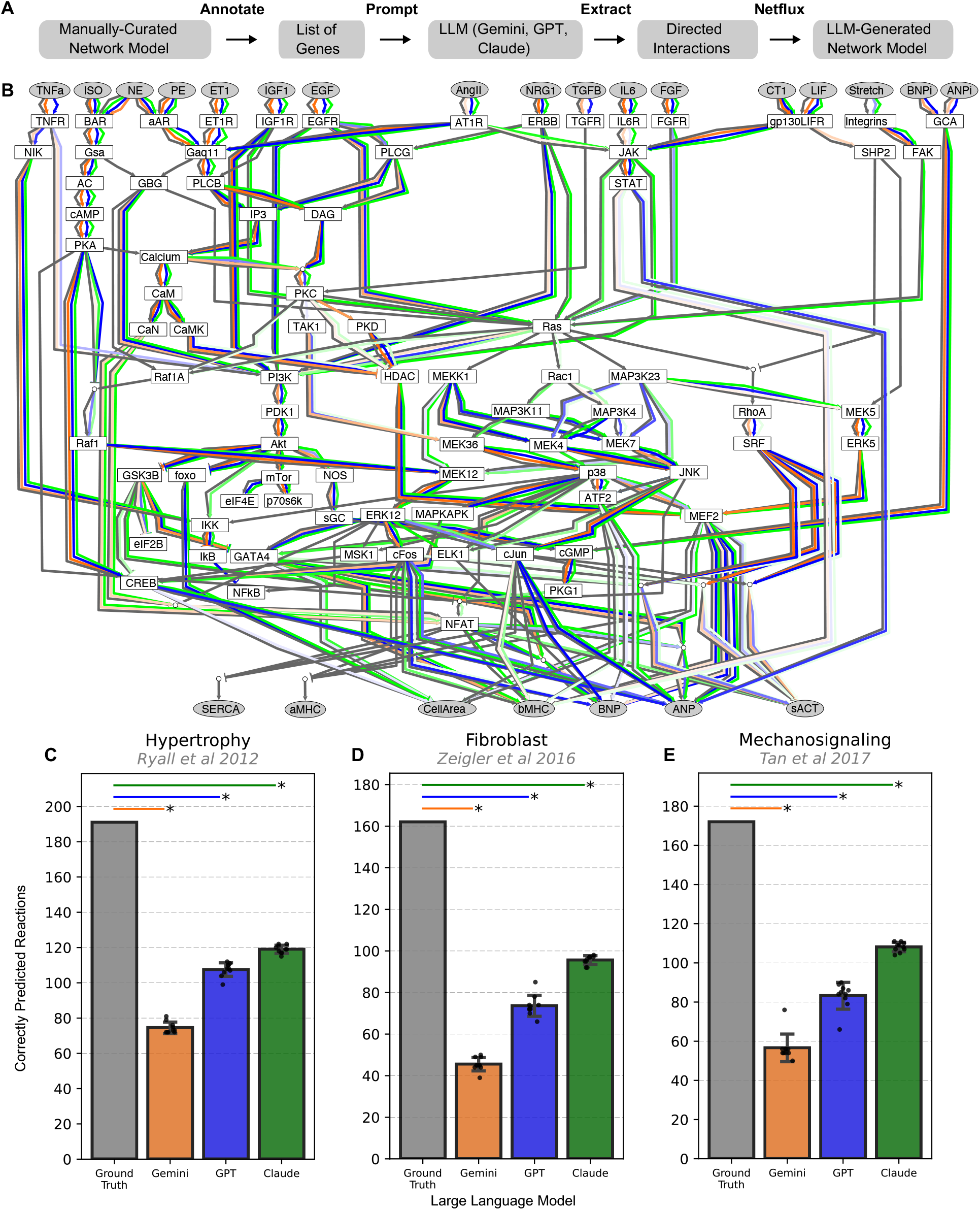
Signaling networks generated by general-purpose large language models. A) Schematic of the pipeline for LLM-generated models of signaling networks. B) Network reactions recalled by three large language models (Gemini 3.0, orange; GPT5.2, blue; Claude4.6, green) compared with a “Ground Truth” literature-curated and validated cardiomyocyte hypertrophy signaling network^9^ (gray reactions). Edge color intensities indicate how frequently each reaction was predicted among the replicates (*n* = 10). LLM networks were generated using iterative prompts based on the gene set of the Ground Truth hypertrophy network. C–E) Summary of reaction recall for three literature-curated signaling networks (C, hypertrophy^9^; D, fibroblast^10^; and xsE, mechanosignaling^11^) by Gemini, GPT, and Claude. * Indicates p < 10^−9^ in one-sample *t-*test between LLM-generated replicates and the ground truth network.

Leveraging this pipeline, we tested the extent to which three general-purpose LLMs (GPT-5.2 Pro, OpenAI; Gemini 3.0 Pro-Preview, Google; and Claude Opus 4.6, Anthropic) could reconstruct the structure of the manually curated cardiomyocyte hypertrophy signaling network^9^, based on its gene annotations alone.

All three LLMs captured aspects of this manually curated network, but with varying coverage depending on the region of the network and the specific LLM (**Figure 1B**). In particular, GPT 5.2 Pro, Gemini 3.0 Pro-preview, and Claude Opus 4.6 all performed relatively well in reconstructing upstream ligand-receptor interactions (e.g. EGF to EGFR) and well-established second-messenger cascades such as the β-adrenergic receptor–cAMP–CREB or phospholipase C–diacylglycerol–PKC pathways. They also accurately predicted the structure of highly conserved signaling axes such as PI3K-Akt-mTor and calcium-calmodulin-calcineurin-NFAT. In contrast, the three LLMs were less able to reconstruct regulation of downstream transcription factors (e.g. cFOS, NFAT, cJUN, MEF2) and gene expression (SERCA, αMHC, βMHC, natriuretic peptides ANP/BNP) or cell area / hypertrophy. The lower reconstruction accuracy of downstream signaling may arise from cardiomyocyte-specific reactions. Overall, the three LLMs reconstructed 37.70-63.87% of network reactions in the cardiomyocyte hypertrophy network (**Figure 1C**).

To what extent is the limited recall of LLM-generated networks specific to cardiomyocyte hypertrophy? To address this question, we used the same reasoning models to reconstruct network models of fibroblast activation^10^ (91 nodes, 134 reactions) and mechanosignaling^11^ (94 nodes, 125 reactions). Like the hypertrophy model, these networks were previously manually reconstructed from the literature, translated into predictive logic-based models, and had their predictions validated against substantial experiments from the literature not used to develop the model. Consistent with the hypertrophy network, LLM-generated models of fibroblast activation and mechanosignaling reconstructed 24.07-60.49% and 29.07-64.53% of their manually curated counterparts (**Figure 1D and E**).

We then asked whether the regional coverage of LLM-generated networks of fibroblasts and mechanosignaling mirrors that of cardiomyocytes. Indeed, the LLMs consistently reconstructed ligand-receptor interactions and highly conserved pathways such as cAMP and Ras/Raf/ERK, but did not consistently recall AT1R signaling, PDGFR-abl signaling, and downstream regulation of transcription factors, genes, and extracellular matrix (e.g. MMP-collagen subnetwork) (**Supplementary Figure 1**). Unlike hypertrophy and fibroblast networks, mechanosignaling is mediated by stretch-dependent proteins that are relatively less characterized. Indeed, the LLM-generated networks for mechanosignaling did not adequately enumerate direct mechanosensory reactions and also had limited coverage of gene regulation (**Supplementary Figure 2**). Still, core conserved pathways for calcium regulation, regulation of protein synthesis, and MAPK signaling were reconstructed robustly.

Models of signaling networks are useful not just for their structure, but also for simulating the response to new perturbations. Prior to benchmarking, the hypertrophy, fibrosis, and mechanosignaling networks were automatically translated into logic-based differential equation models using the Netflux framework^12,13^. This allows network-wide prediction of dynamics in response to desired perturbations and does not require prior knowledge of rate constants or concentrations^13^. The predictions of these three logic-based models have been rigorously validated against experimental evidence not used to develop the model^9–11,14^. Using the same Netflux approach, we generated logic-based models based on all of the LLM-generated network structures shown in **Figure 1**.

Returning our focus to the hypertrophy network, we examined the ability of LLM-generated networks to predict the classic “fetal gene program” gene expression signature in cardiomyocytes. The manually curated model accurately predicts that AngII and/or ISO increase the expression of ANP, BNP, GATA4, βMHC, sACT, and increase cell area/size, while decreasing expression of SERCA and αMHC. In contrast, the LLM-generated network models produce limited aspects of this signature, with the best performance in predicting ISO-dependent ANP in the GPT version (**Figure 2A**). We then tested whether this validation accuracy scales across all available manually curated experiments that cover a wider array of perturbations and measured outputs. Overall, for 114 validations where the manually curated network has a functional accuracy of 94.74%, LLM-generated networks have an accuracy of 6.14-33.33% (**Figure 2B**). We then repeated this systematic experimental validation in comparison to literature-based models of the fibroblast (82% validation against independent literature) and mechanosignaling (78% validation). LLM-generated models of the fibroblast (**Figure 2C**) and mechanosignaling (**Figure 2D**) networks exhibited validation accuracies similar to the hypertrophy model: 16.87-21.69% and 5.81-14.53%, respectively. Across all three networks, validation accuracy of LLM-generated models for perturbation responses is lower than their accuracy for reconstructing network structure.

**Figure 2:**
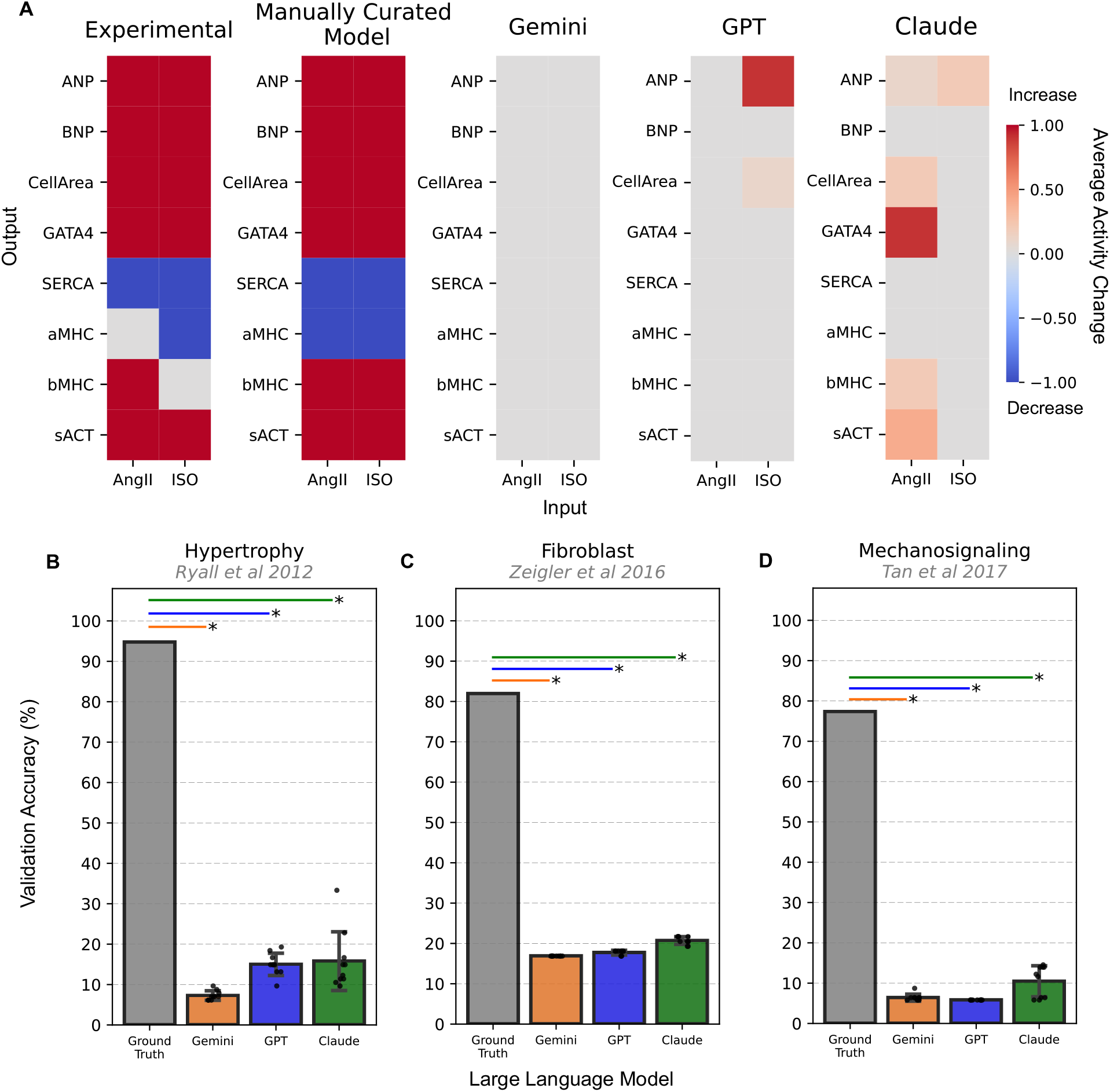
Experimental validation of perturbation responses predicted by LLM-generated signaling network models. A) Representative validations of network models generated by manual curation or by LLMs (Gemini, GPT, Claude), in comparison to experiments in conditions of Angiotensin II (AngII) or isoproterenol (ISO) from the literature^9^. B–D) Summary of systematic validations of manually curated (Ground Truth) and LLM-generated reconstructions of hypertrophy, fibroblast, and mechanosignaling network models against perturbation experiments from the literature (n = 114, 83, and 171 experiments, respectively). * Indicates p < 10^−11^ in one-sample T test between LLM-generated model validation scores (n = 10 replicates) and ground truth model validation accuracy.

It is important to note that the functional validation simulations used LLM-generated signaling network models after pruning to the subset of reactions contained in the ground truth networks. This enabled direct mapping of LLM-predicted interactions onto the “AND” formalism used in the logic-based network models, but also excluded predicted biological interactions that were generated by the LLMs but absent from the curated networks, which we tentatively classify as false positives (LLM^+^, ground-truth^−^). When this constraint was removed and the full set of LLM-predicted interactions was used for functional validation, model performance did not substantially improve, except for the fibroblast network (**Supplementary Figure 6A-C**). Additionally, the average size of the unconstrained LLM-generated networks was substantially larger than the manually curated network (**Supplementary Figure 6D**).

Additional performance metrics, including confusion matrices and calibration analyses for each signaling network, are provided in **Supplementary Figures 3–5**. Notably, Gemini generally exhibited substantially higher precision than Claude or GPT in predicting signaling network reactions. F1 scores were relatively consistent among the evaluated LLMs.

Reactions generated by the LLMs but absent from the ground-truth signaling networks were classified as “false positives” for the purpose of calculating performance metrics. This classification should be interpreted cautiously, as some LLM-predicted/ground-truth-negative reactions may represent true biomolecular interactions or regulatory relationships that were not captured in the manually curated reference models. To assess the impact of retaining these additional predictions, we evaluated the functional performance of the full LLM-generated signaling networks without pruning LLM-predicted/ground-truth-negative reactions. This analysis revealed only marginal improvements, and in some cases reduced accuracy, for the hypertrophy and mechanosignaling networks (**Supplementary Figure 6 A and C**). In contrast, the unrefined fibroblast network models exhibited substantially greater functional validation performance than the pruned reconstructions, which retained only LLM-predicted/ground-truth-positive reactions (**Supplementary Figure 6B**). The Claude- and GPT-generated signaling networks contained more connections than the corresponding manually curated networks – this increased network density may have contributed to their improved functional performance within the fibroblast context (**Supplementary Figure 6D**).

Expanding the scope of our LLM-benchmarking, we asked how effectively these models could reconstruct a metabolic network of core *Escherichia coli* (*E. coli*) metabolism^15^. The reconstruction was guided by the genes represented in the gene– protein–reaction (GPR) associations of the reference network. We adapted our prompting approach to generate the information needed to define metabolic reactions, including substrates, products, stoichiometry, and reversibility. These outputs were used to assemble LLM-generated metabolic network models for comparison against the manually curated reference network, as described in the **Methods**.

There is notable heterogeneity in the regional coverage of this metabolic network by the different LLMs. Generally, both Claude and Gemini were able to accurately reconstruct the glycolysis and tricarboxylic acid cycle (TCA) while GPT failed to capture a few specific reactions in each process including the phosphoglycerate mutase and kinase (PGM and PGK respectively). Additionally, GPT failed to reconstruct the lactate dehydrogenase and lactate transport reactions. Gemini and Claude performed poorly in recreating the malate and fumarate transport reactions feeding into the TCA cycle. Overall, the LLM-generated metabolic network models demonstrated markedly greater coverage of the ground-truth core *E. coli* metabolic network than the predicted signaling networks achieved for their respective ground-truth networks. Specifically, the LLM-generated models reconstructed 63.63–90.90% of reactions within the manually curated core *E. coli* network (**Figure 3C**).

Despite substantial improvement in coverage of the ground truth reactions, the LLM-generated metabolic networks models largely underperformed in the substrate utilization analysis. Indeed, the majority of the networks yielded infeasible flux solutions or predicted no growth on all of the substrates (**Figure 3D**). Interestingly, a sparse subset of the Gemini- and Claude-generated models achieved strong substrate utilization prediction performance, with accuracies approaching the score of the ground-truth model (**Figure 3E**).

**Figure 3:**
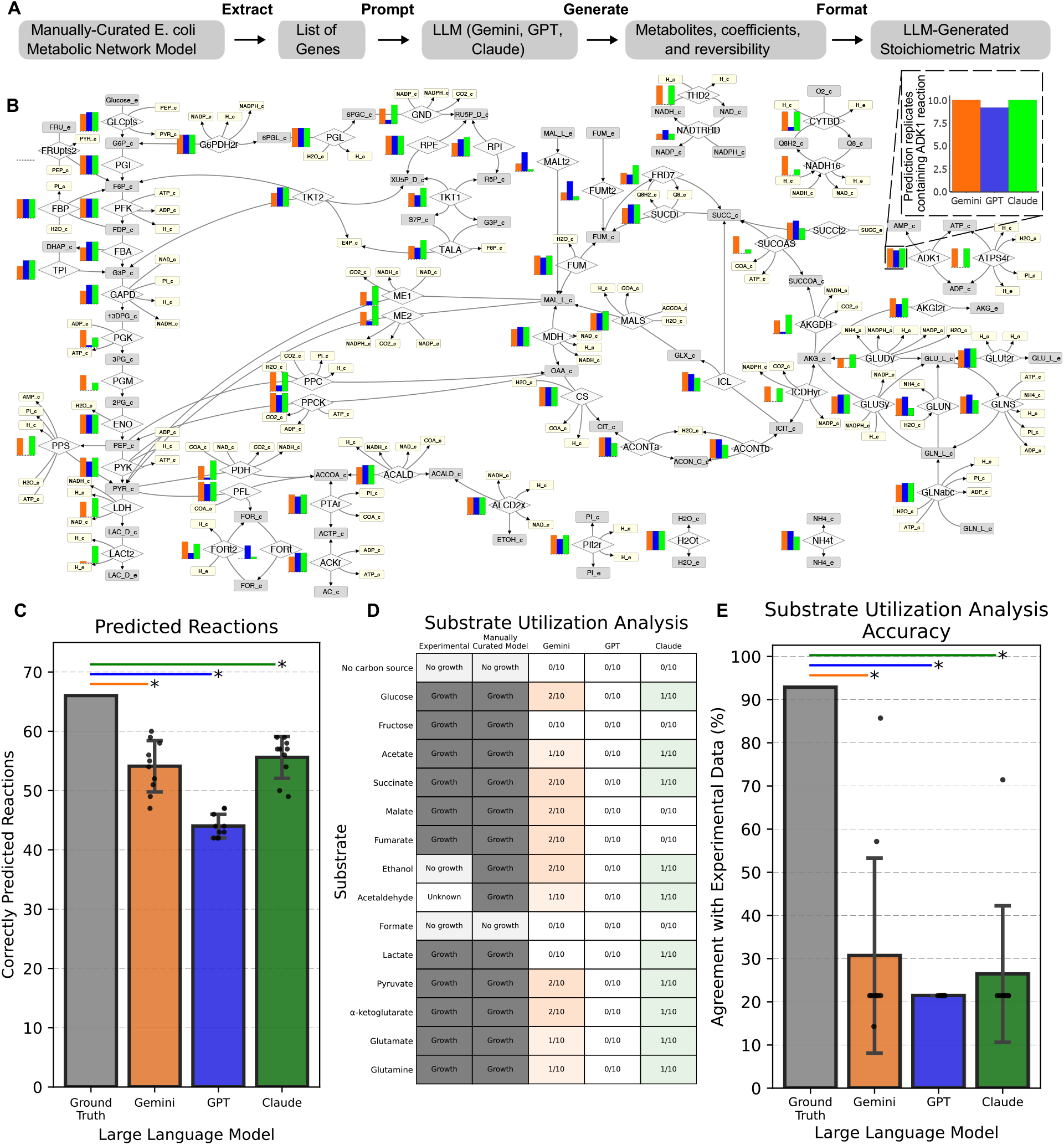
Metabolic network models generated by general-purpose LLMs and substrate utilization analysis. A) Schematic of pipeline for LLM-generated models of the core *E. coli* metabolic network. B) Network reactions recalled by three large language models (Gemini, orange; GPT, blue; Claude, green) compared with a “Ground Truth” manually curated metabolic network. Bar charts illustrate how frequently each reaction was predicted by the different LLMs in the replicates (n = 10). Inset shows zoomed-in model coverage of the ADK1 reaction. LLM networks were generated using iterative prompts based on the GPR gene list of the core *E. coli* metabolic network. C) Summary of reaction recall accuracy by Gemini, GPT, and Claude. D) Heatmap illustrating substrate utilization predictions of the ground truth model to and LLM-generated models compared to experimental data. Color intensity within the LLM columns indicates frequency of growth prediction amongst the 10 replicates. E) Summary of model performance compared to the experimental data across carbon sources. * Indicates p < 10^−4^ in one-sample T test between LLM-generated replicates (n = 10) and the ground truth network. Within the network visualization, diamonds indicate reactions while rectangles indicate metabolites.

In conclusion, we developed pipelines that use general-purpose LLMs to generate predictive models of signaling and metabolic networks. We benchmarked the signaling network pipeline across three LLMs against large-scale, manually curated, and experimentally validated network models. Across cardiac hypertrophy, fibroblast, and mechano-signaling networks, LLM-generated models exhibit 24.07-64.53% reconstruction of reactions from the manually curated signaling models. Reconstruction accuracy is high for ligand-receptor interactions and well-conserved pathways, but lower for cell-specific gene regulation or biophysical regulation.

Similarly, we benchmarked the LLMs’ performance in generating a metabolic network using experimental data and a manually curated reference model. Although the LLMs achieved greater structural recovery of the core *E. coli* metabolic network relative to the signaling networks, these reconstructions did not translate into robust functional performance as assessed by substrate utilization analysis. This discrepancy may reflect gaps in key reconstructed metabolic pathways and aligns with findings from the signaling network models, where incomplete reconstruction of critical pathways likely limited the accuracy of simulated perturbation responses. The ability of LLMs to reconstruct core *E. coli* metabolism may reflect the depth of existing literature on this model system, and similar approaches may perform worse for metabolic networks that are less extensively characterized.

This benchmarking does not yet test the ability of LLMs to reconstruct aspects of signaling that are not represented in literature-curated networks, such as isoform-specific regulation^16^ or liquid-liquid phase separation^17^. Such mechanisms are not yet adequately characterized for global inclusion in either literature-curated or LLM-generated models.

Future efforts could build on the pipelines developed in this work. Improving the performance of LLM-generated network models may require hybrid manual–LLM workflows or specialized AI models tailored to biological network reconstruction. Another promising direction would be to implement an agentic framework in which some agents propose plausible signaling reactions between genes, while others serve in a reviewer role to critique and approve candidate reactions. Such structured critique paradigms have shown promise in AI-oriented research contexts^18^. As LLMs and other AI models improve in their ability to synthesize and connect biological knowledge derived from experimental studies, developing rigorous platforms for organizing and evaluating this information will be increasingly important.

## Methods

### Prior knowledge models of signaling networks

We benchmarked LLM-generated network models using three large-scale network models based on manual curation of directed interactions using the literature^9–11^. These models contain a range of biological processes including ligand-receptor interactions, phosphorylation, gene regulation, and cellular phenotypes. Each node is annotated with corresponding genes. The models were previously implemented as systems of logic-based differential equations^13^, enabling validation of predictions to new perturbations with accuracies of 94.74% (hypertrophy), 81.93% (fibroblast), and 77.33% (mechanosignaling)^9–11^. Validation simulations were performed with parameter sets from each manually curated signaling network model’s original publication^9–11^.

### Querying large language models to reconstruct signaling networks

We evaluated three widely used, general-purpose large language models (LLMs): GPT-5.2 Pro (OpenAI), Gemini 3.0 Pro-preview (Google), and Claude Opus 4.6 (Anthropic). For each model, prompting was performed through the corresponding application programming interface (API) using structured outputs to standardize response formatting across models. Temperature was set to 1.0, the maximum output length was set to 10,000 tokens, and 10 replicate outputs were generated for each LLM–signaling network pair, yielding 90 total outputs. The prompts consisted of three steps:

1. The list of gene symbols, along with the description: “List of genes and other signaling nodes:”.
2. Description of the desired outputs: “For the first {batch_size} entries in this list of genes, proteins, and other signaling nodes from a {desc} signaling network, please provide more than 0 but fewer than {max_connections} direct interactions with other nodes from the list supported by available literature. Return plausible interactions between these nodes in JSON format. Each interaction should have: source_node, target_node, and action (“stimulation” or “inhibition”).” Batch size was set at 20 genes, max connections set to the maximum number of connections exhibited in that parent network, and phenotype set at “cardiac hypertrophy”, “fibroblast”, or “mechanosignaling” corresponding to the three networks used for benchmarking.
3. Instructions to repeat the process until reaching the end of the gene list: “That looks great! Please do the same operation for the next {len(chunk)} nodes! Thank you”

### Storing LLM-output in structured format and converting to logic-based models

Structured LLM outputs for the signaling networks were stored in JavaScript Object Notation (JSON) as interaction lists with three standardized fields: source_node, target_node, and action. The action field encoded the predicted regulatory relationship as either stimulation or inhibition. JSON interaction lists were converted into Netflux format for automated conversion into logic-based ordinary differential equation models^12,13^. These logic-based models were subsequently used for functional validation simulations as shown in **Figure 2**. In **Figure 1**, we evaluated restrictive reconstructions of the predicted signaling networks by retaining only LLM-predicted reactions that were also present in the corresponding ground-truth network. This approach excluded LLM-predicted/ground-truth-negative reactions, limiting the functional evaluation to true-positive reactions. This approach is in contrast with that of **Supplementary Figure 6** which illustrates functional evaluation of unrefined, full LLM-generated networks.

### Constructing confusion matrices using LLM-generated signaling network models

For each predicted signaling network model, we calculated sensitivity, specificity, precision, negative predictive value (NPV), accuracy, and F1 score relative to the corresponding ground-truth network. These metrics were summarized as confusion matrices for the hypertrophy, fibroblast, and mechanosignaling benchmarks (**Supplementary Figures 3–5**). Actual positives were defined as node-to-node connections present in the ground-truth network, whereas actual negatives were defined as all possible node-to-node connections among ground-truth nodes that were absent from the manually curated model.

Because the number of absent node-to-node connections greatly exceeded the number of curated reactions, these classification tasks were characterized by substantial class imbalance. For example, the hypertrophy network contained 191 curated connections among 106 nodes, yielding 11,236 possible directed node-to-node connections and 11,045 actual negatives. Consequently, true negatives dominated the confusion matrices, making specificity, accuracy, and NPV less informative than precision, sensitivity, and F1 score.

This effect was illustrated using null predictors, defined as hypothetical models that predicted no connections. These null predictors had sensitivity values of 0 and specificity values of 1, while still achieving high accuracy and NPV because of the large number of true negatives.

### Calibration analyses and predicted model reaction unions

Calibration analyses were enabled by the generation of 10 replicates per LLM per signaling network model. The Scikit Learn calibration curve package was utilized to visualize the results of this analysis^19^. The Venn Diagrams contained in Supplementary Figures 3-5 illustrate the total reaction populations predicted by the LLMs (across all replicates) and their overlap with the ground truth model reaction set.

### Querying large language models to reconstruct a metabolic network

The core *E. coli* model contains only central carbon metabolism^15^. This small-scale model (72 metabolites, 95 reactions, and 137 genes) contains reactions of glycolysis, the pentose phosphate shunt, the tricarboxylic acid cycle, the glyoxylate cycle, gluconeogenesis, anaplerotic reactions, the electron transport chain, fermentation, the transfer of reducing equivalents, and nitrogen metabolism. The model was downloaded from the BiGG Database (ID: e_coli_core), and the associated gene symbols were extracted^20^. Similar to the signaling network strategy, these genes were passed to the LLM using a sliding window approach with structured outputs (JSON format) and 10 replicates were generated per LLM. The LLM versions, temperature, maximum tokens, and other prompting parameters were identical to those used in generating signaling networks. The specific prompt is included below:

> *“For the first {batch_size} entries in this list of genes from an E. coli metabolic network, please return a structured list of metabolic reactions using the provided JSON schema:*
>
> *Each interaction should have: a reaction_id, substrates (list of metabolite and coefficient pairs in BiGG format with compartment suffixes ‘_c’ or ‘_e’), products (list of metabolite and coefficient pairs in BiGG format with compartment suffixes ‘_c’ or ‘_e’), and reversible (boolean). If a gene catalyzes multiple reactions, please return each reaction separately*.
>
> *[iteration completed]*
>
> *That looks great! Please do the same operation for the next {len(chunk)} genes! Thank you”*

Out of the 95 reactions in this core network, 26 are not associated with specific gene symbols and constitute transport, sink, demand, exchange, or spontaneous reactions. Additionally, three transport reactions (oxygen, carbon dioxide, and acetaldehyde) are associated with a placeholder gene ‘s0001’. Therefore, reaction-recovery analysis was performed using the 66 gene-associated reactions as the reference set.

Precise BiGG notation was requested for the predicted metabolite sets to facilitate mapping back to the ground truth model. Because the JSON schema stored substrate and product coefficients as positive magnitudes, substrate coefficients were multiplied by −1 when constructing stoichiometric matrices. Metabolite IDs were mapped to BiGG metabolite IDs using case-insensitive matching. This enabled generation of stoichiometric matrices for each replicate with matched metabolite IDs to the ground truth as rows and arbitrary reaction IDs (generated by the LLM but not formatted to BiGG notation) as columns. To identify the reactions predicted by the models, we leveraged the metabolite IDs and coefficients to perform vector comparison to the ground truth model’s columns (curated reactions). This vector-based matching was performed for all non-zero coefficient columns in the generated stoichiometric matrices. If the predicted model included non-zero coefficients for a set of metabolites that precisely matched a reaction column in the ground truth matrix (set of non-zero coefficients), that reaction was designated as ‘recovered.’ Metabolites that could not be mapped to BiGG IDs were excluded from strict matching and reversibility annotations were not considered for the recovery analysis.

### Substrate utilization analysis using LLM-generated metabolic network models

To perform substrate utilization analysis, we leveraged an analogous approach to the signaling network validations. The predicted reactions that matched ground-truth reactions (i.e. ‘true-positive’ reactions) were combined into a reconstructed metabolic network model that enabled flux balance analysis (FBA). FBA, which underlies substrate utilization analysis, requires a sufficiently connected metabolic network to admit feasible flux solutions; thus, several of these predicted networks yielded infeasible solutions for particular substrates. Reconstructed networks were supplemented with exchange, demand, sink, biomass, ATP maintenance, and other non-gene-associated reactions to enable full FBA. The LLM-generated reversibility Boolean annotations were used to assign reaction bounds. Reactions with reversible = TRUE were assigned bounds of [-1000, 1000], whereas irreversible reactions were assigned bounds of [0, 1000] in agreement with BiGG conventions.

All of the carbon sources for the model were tested including glucose, fructose, acetate, succinate, malate, fumarate, ethanol, acetaldehyde, formate, lactate, pyruvate, alpha-ketoglutarate, glutamate, and glutamine. For each substrate condition, the corresponding exchange reaction lower bound was set to −10, while other tested carbon-source exchange reactions were closed. All other medium components were set according to the default medium in the BiGG model. The minimum threshold to define growth was 10^−6^. FBA was performed using COBRApy v0.30.0 and the biomass reaction was used as the objective^21^. Experimental data was taken from the EcoCyc database’s list of growth-media for *E. coli* substrain MG1655^22^. This database contains consensus experimental findings (‘growth’, ‘weak growth’, ‘no growth’, or ‘inconclusive’) for *E. coli MG1655’s* growth at 37° C under aerobic conditions using isolated carbon sources. Two supplementary publications provided experimental evidence that unengineered *E. coli* cannot grow when ethanol is supplied as the sole carbon source^23,24^.

The code and LLM-generated predictions in JSON format are freely available on GitHub at: https://github.com/saucermanlab/LLM-network-generation

## Supplementary Figure Legends

**Supplementary Figure 1. Visualization of LLM-generated fibroblast signaling networks, as recalled by three general-purpose large language models**. Network reactions recalled by three large language models compared with a “Ground Truth” literature-curated and validated fibroblast signaling network (gray reactions). LLM-generated networks used prompts based on the gene set of the Ground Truth fibroblast network. This visualization corresponds to the analyses in **Figure 1D**.

**Supplementary Figure 2. Visualization of LLM-generated mechanosignaling networks, as recalled by three general-purpose large language models**. Network reactions recalled by three large language models (Gemini3.0, orange; ChatGPT-5.2, blue; Claude4.6, green) compared with a “Ground Truth” literature-curated and validated mechanosignaling network (gray reactions). LLM-generated networks used prompts based on the gene set of the Ground Truth mechanosignaling network. This visualization corresponds to the analyses in **Figure 1E**.

**Supplementary Figure 3. Confusion matrices, calibration analysis, and intersection of predicted reactions across replicates for the LLM-generated hypertrophy networks**. A) Confusion matrices summarizing reaction-level prediction performance for each LLM and a null predictor (hypothetical network with no connections) relative to the manually curated hypertrophy network. Actual positives were defined as reactions present in the manually curated network, whereas actual negatives were defined as possible node-to-node reactions absent from the manually curated network. B) Calibration analyses were performed using replicate predictions for each LLM (*n* = 10), with confidence defined as the fraction of replicates in which a reaction was predicted. C) Venn diagram showing the overlap between the manually curated ground-truth reactions and the union of reactions predicted by at least one replicate from each LLM (confidence ≥ 0.1). TP, true positive; FP, false positive; TN, true negative; FN, false negative; NPV, negative predictive value; F1, F1 score.

**Supplementary Figure 4. Confusion matrices, calibration analysis, and intersection of predicted reactions across replicates for the LLM-generated fibroblast networks**. A) Confusion matrices summarizing reaction-level prediction performance for each LLM and a null predictor relative to the manually curated fibroblast network. Actual positives were defined as reactions present in the manually curated network, whereas actual negatives were defined as possible node-to-node reactions absent from the manually curated network. B) Calibration analyses were performed using replicate predictions for each LLM (*n* = 10), with confidence defined as the fraction of replicates in which a reaction was predicted. C) Venn diagram showing the overlap between the manually curated ground-truth reactions and the union of reactions predicted by at least one replicate from each LLM (confidence ≥ 0.1). TP, true positive; FP, false positive; TN, true negative; FN, false negative; NPV, negative predictive value; F1, F1 score.

**Supplementary Figure 5. Confusion matrices, calibration analysis, and intersection of predicted reactions across replicates for the LLM-generated mechanosignaling networks**. A) Confusion matrices summarizing reaction-level prediction performance for each LLM and a null predictor relative to the manually curated mechanosignaling network. Actual positives were defined as reactions present in the manually curated network, whereas actual negatives were defined as possible node-to-node reactions absent from the manually curated network. B) Calibration analyses were performed using replicate predictions for each LLM (*n* = 10), with confidence defined as the fraction of replicates in which a reaction was predicted. C) Venn diagram showing the overlap between the manually curated ground-truth reactions and the union of reactions predicted by at least one replicate from each LLM (confidence ≥ 0.1). TP, true positive; FP, false positive; TN, true negative; FN, false negative; NPV, negative predictive value; F1, F1 score.

**Supplementary Figure 6. Experimental validation of perturbation responses predicted by full LLM-generated signaling network models**. A–C) Systematic validation of manually curated ground-truth models and full, unrefined LLM-generated models for the hypertrophy, fibroblast, and mechanosignaling networks against perturbation experiments from the literature (n = 114, 83, and 171 experiments, respectively). Asterisks indicate p < 1 × 10^−7^ by one-sample t-test comparing LLM-generated model validation scores across replicates (n = 10 per LLM) against the corresponding ground-truth model validation accuracy. D) Average reaction counts in the full LLM-generated network models (n = 10 per network per LLM) compared with corresponding manually curated networks.

