## Supplementary Figures for "Benchmarking biochemical networks generated by large language models"

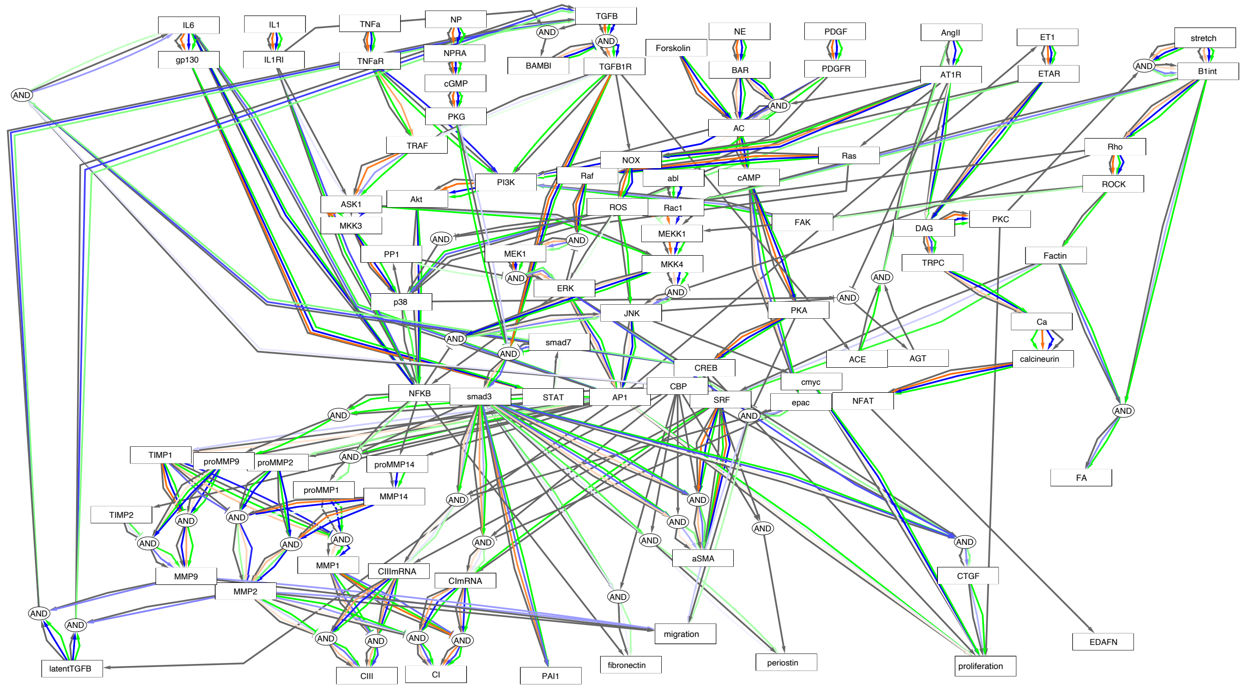


**Supplementary Figure 1. Visualization of LLM-generated fibroblast signaling networks, as recalled by three general-purpose large language models.** Network reactions recalled by three large language models compared with a "Ground Truth" literature-curated and validated fibroblast signaling network (gray reactions). LLM-generated networks used prompts based on the gene set of the Ground Truth fibroblast network. This visualization corresponds to the analyses in **Figure 1D**.

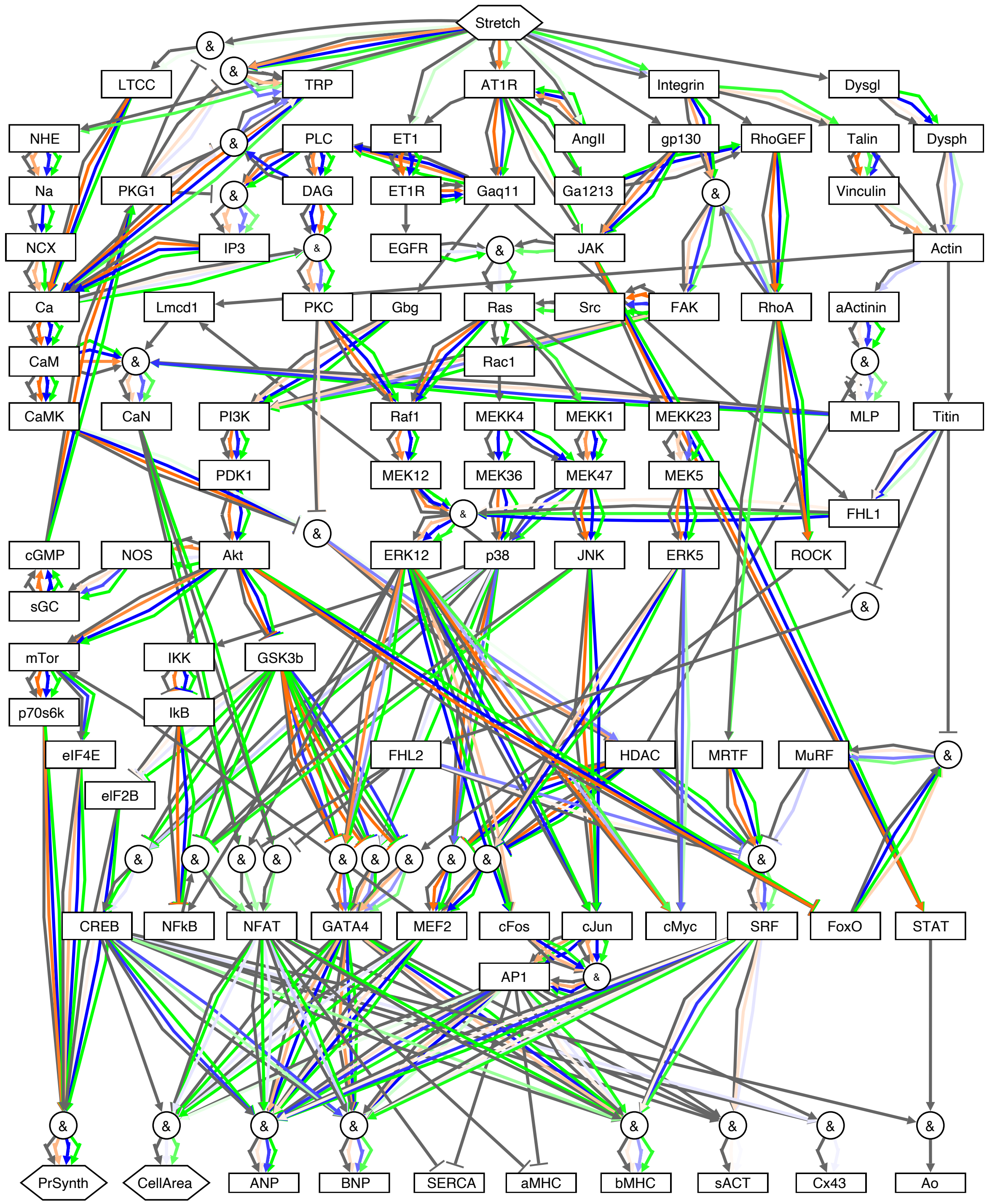


**Supplementary Figure 2. Visualization of LLM-generated mechanosignaling networks, as recalled by three general-purpose large language models.** Network reactions recalled by three large language models (Gemini3.0, orange; ChatGPT-5.2, blue; Claude4.6, green) compared with a "Ground Truth" literature-curated and validated mechanosignaling network (gray reactions). LLM-generated networks used prompts based on the gene set of the Ground Truth mechanosignaling network. This visualization corresponds to the analyses in **Figure 1E**.

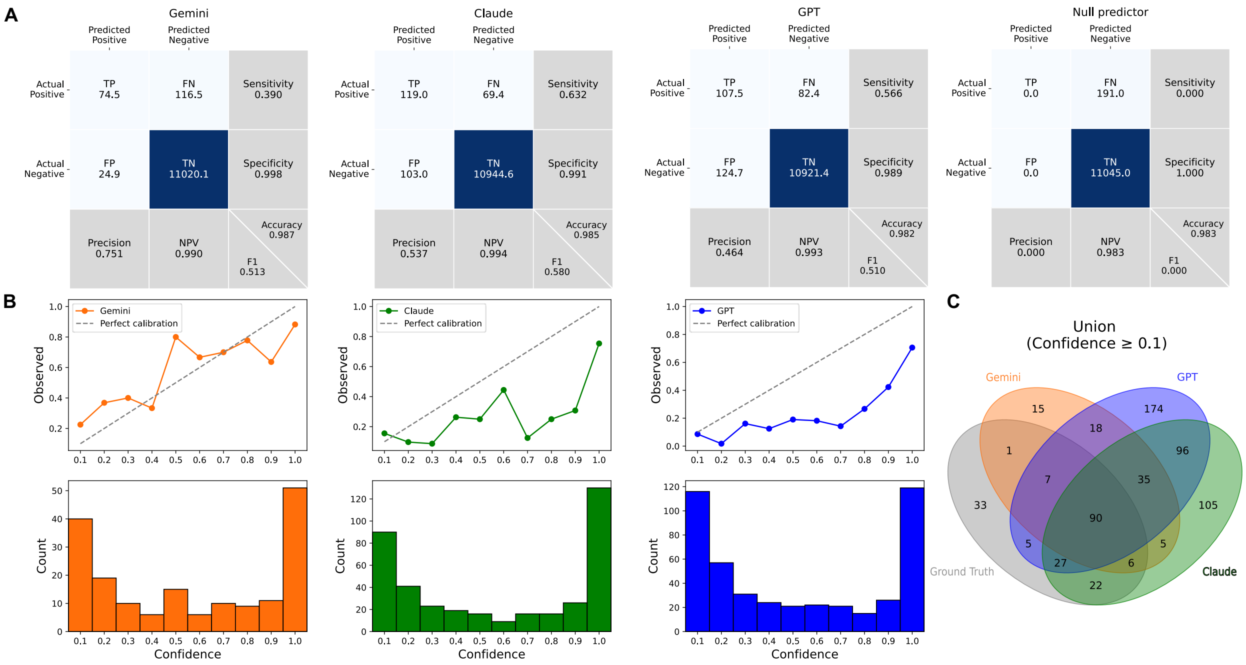


**Supplementary Figure 3. Confusion matrices, calibration analysis, and intersection of predicted reactions across replicates for the LLM-generated hypertrophy networks.** A) Confusion matrices summarizing reaction-level prediction performance for each LLM and a null predictor (hypothetical network with no connections) relative to the manually curated hypertrophy network. Actual positives were defined as reactions present in the manually curated network, whereas actual negatives were defined as possible node-to-node reactions absent from the manually curated network. B) Calibration analyses were performed using replicate predictions for each LLM ($n$ = 10), with confidence defined as the fraction of replicates in which a reaction was predicted. C) Venn diagram showing the overlap between the manually curated ground-truth reactions and the union of reactions predicted by at least one replicate from each LLM (confidence ≥ 0.1). TP, true positive; FP, false positive; TN, true negative; FN, false negative; NPV, negative predictive value; F1, F1 score.

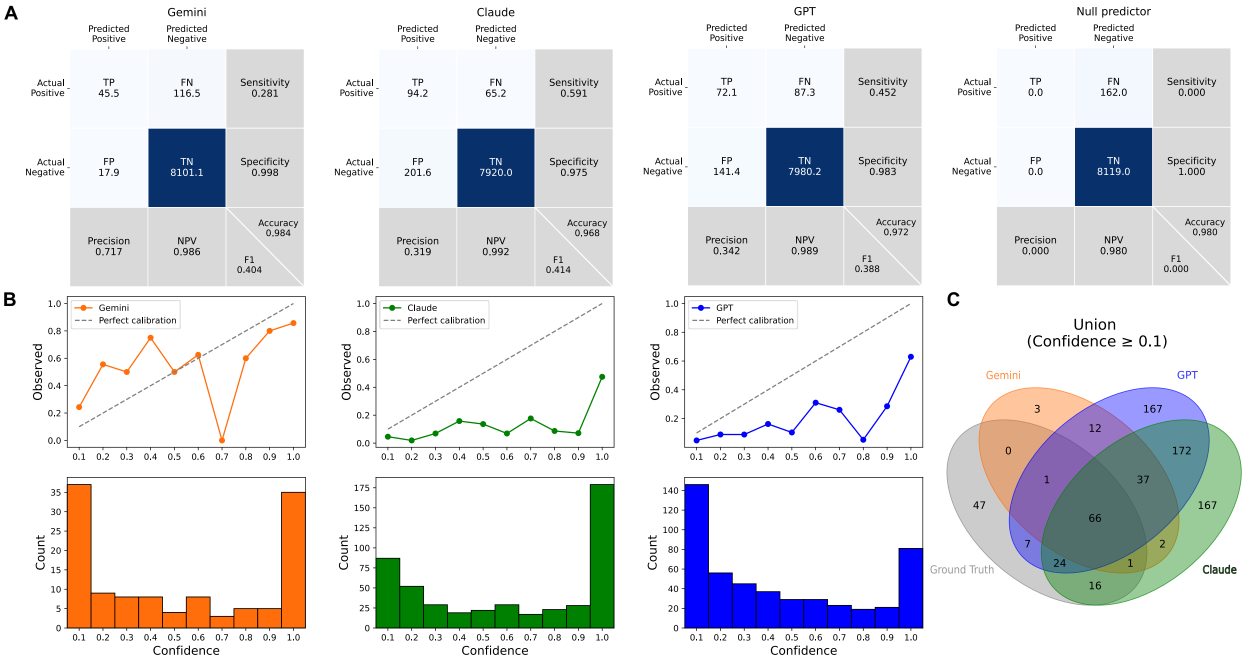


**Supplementary Figure 4. Confusion matrices, calibration analysis, and intersection of predicted reactions across replicates for the LLM-generated fibroblast networks.** A) Confusion matrices summarizing reaction-level prediction performance for each LLM and a null predictor relative to the manually curated fibroblast network. Actual positives were defined as reactions present in the manually curated network, whereas actual negatives were defined as possible node-to-node reactions absent from the manually curated network. B) Calibration analyses were performed using replicate predictions for each LLM (*n* = 10), with confidence defined as the fraction of replicates in which a reaction was predicted. C) Venn diagram showing the overlap between the manually curated ground-truth reactions and the union of reactions predicted by at least one replicate from each LLM (confidence ≥ 0.1). TP, true positive; FP, false positive; TN, true negative; FN, false negative; NPV, negative predictive value; F1, F1 score.

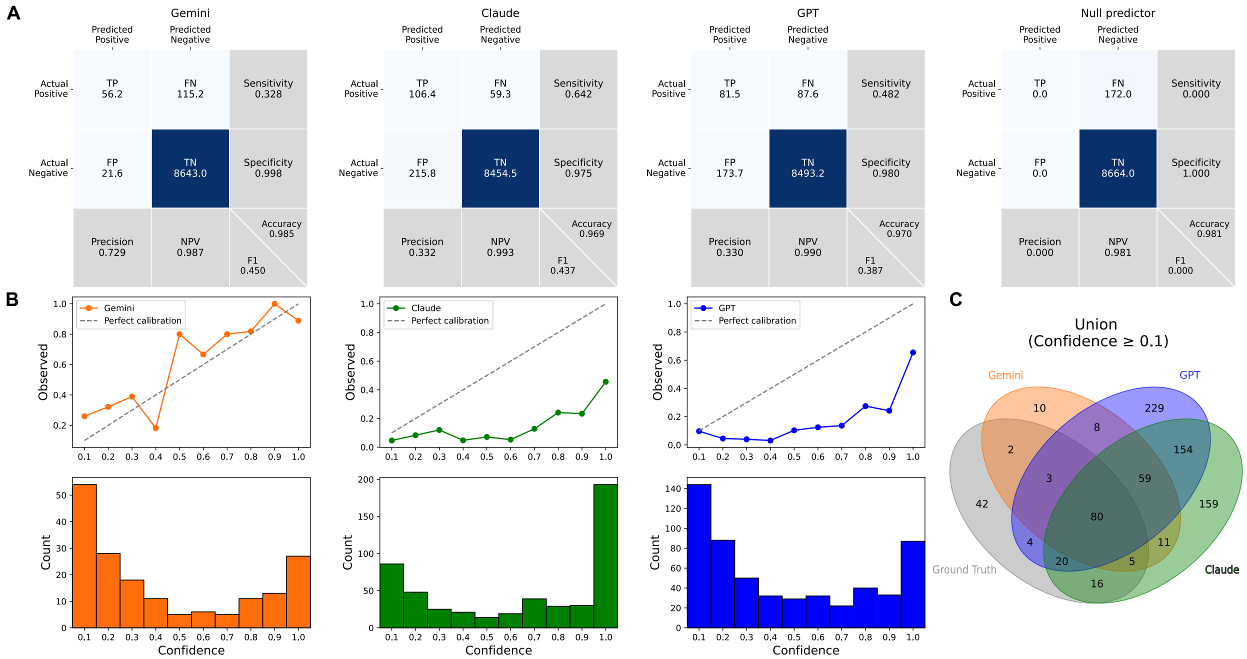


**Supplementary Figure 5. Confusion matrices, calibration analysis, and intersection of predicted reactions across replicates for the LLM-generated mechanosignaling networks.** A) Confusion matrices summarizing reaction-level prediction performance for each LLM and a null predictor relative to the manually curated mechanosignaling network. Actual positives were defined as reactions present in the manually curated network, whereas actual negatives were defined as possible node-to-node reactions absent from the manually curated network. B) Calibration analyses were performed using replicate predictions for each LLM ($n$ = 10), with confidence defined as the fraction of replicates in which a reaction was predicted. C) Venn diagram showing the overlap between the manually curated ground-truth reactions and the union of reactions predicted by at least one replicate from each LLM (confidence ≥ 0.1). TP, true positive; FP, false positive; TN, true negative; FN, false negative; NPV, negative predictive value; F1, F1 score.

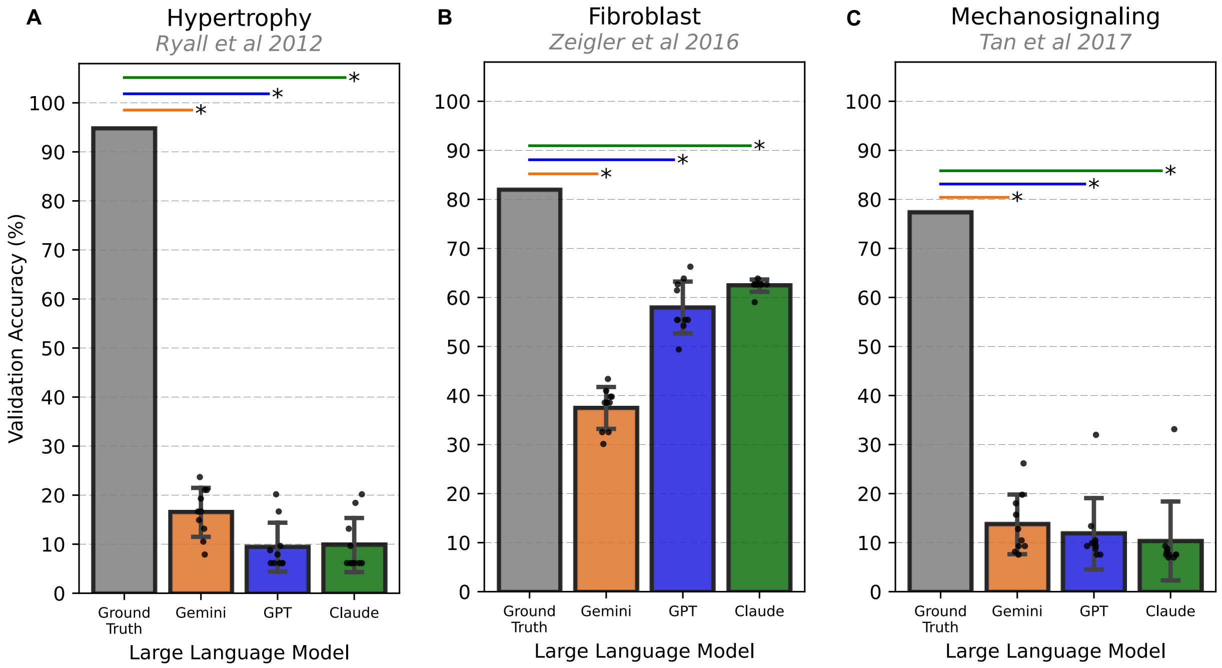


D

| Reaction Counts | Ground Truth | Gemini | GPT | Claude |
|---|---|---|---|---|
| Hypertrophy | 191 | 99.3 | 230.5 | 221.5 |
| Fibroblast | 162 | 62.0 | 214.1 | 294.1 |
| Mechanosignaling | 172 | 77.9 | 256.6 | 323.0 |

**Supplementary Figure 6. Experimental validation of perturbation responses predicted by full LLM-generated signaling network models.** A–C) Systematic validation of manually curated ground-truth models and full, unrefined LLM-generated models for the hypertrophy, fibroblast, and mechanosignaling networks against perturbation experiments from the literature (n = 114, 83, and 171 experiments, respectively). Asterisks indicate $p < 1 \times 10^{-7}$ by one-sample t-test comparing LLM-generated model validation scores across replicates (n = 10 per LLM) against the corresponding ground-truth model validation accuracy. D) Average reaction counts in the full LLM-generated network models (n = 10 per network per LLM) compared with corresponding manually curated networks.
